# Temporal dynamics of metabolic acquisition in grafted engineered human liver tissue

**DOI:** 10.1101/2022.07.21.501040

**Authors:** Chelsea L. Fortin, Tara N. McCray, Sarah H. Saxton, Fredrik Johansson, Christian B. Andino, Jonathan Mene, Yuliang Wang, Kelly R. Stevens

**Affiliations:** Department of Laboratory Medicine & Pathology, University of Washington, Seattle, Washington, USA, 98195; Institute for Stem Cell & Regenerative Medicine, University of Washington, Seattle, Washington, USA, 98109; Department of Bioengineering, University of Washington, Seattle, Washington, USA, 98105; Department of Computer Science & Engineering, University of Washington, Seattle, Washington, USA, 98195

**Keywords:** liver, zonation, metabolism, bioengineering, RNAseq

## Abstract

Liver disease affects millions globally and end-stage liver failure is only cured by organ transplant. Unfortunately, there is a growing shortage of donor organs and disparities in equitable access to transplants among different populations. Less than 10% of global transplantation needs are currently met, highlighting the demand for alternative therapies. Engineered liver tissue grafts that supplement organ function could address these demands. While engineered liver tissues built from human hepatocytes, endothelial cells, and fibroblasts encased in hydrogel have been successfully engrafted in rodent models previously, the extent to which these tissues express human liver metabolic genes and proteins remains unknown. Here, we built engineered human liver tissues and characterized their engraftment, expansion, and metabolic phenotype at sequential stages post-implantation by RNA sequencing, histology, and host serology. Expression of metabolic genes was observed at weeks 1-2, followed by cellular organization into hepatic cords by weeks 4-9.5. Furthermore, grafted engineered tissues exhibited progressive spatially restricted expression of critical functional proteins known to be zonated in the native human liver. To our knowledge, this is the first report of engineered human liver tissue zonation after implantation in vivo, which could have important translational implications for this field.

## 1. Introduction

The liver is the largest internal organ and performs hundreds of essential functions for human health. Liver disease is a significant global health burden, causing over 40,000 annual deaths in the United States alone and more than one million worldwide deaths each year.^[1]^ While liver transplant is curative, less than ten thousand occur per year in the US.^[2]^ There is also a growing shortage of donor organs and disparities in equitable access to transplants amongst non-white, women, uninsured and rural populations.^[3]^ Less than 10% of global transplantation needs are currently met,^[4]^ highlighting the demand for alternative therapies. Engineered liver tissue that supplements organ function could bridge patients to transplant and help alleviate the donor tissue shortage.^[5]^ To reach the clinic, engineered liver tissues would ideally mimic key structural and physiological aspects of the native liver.

The hundreds of functions performed by the native liver can be broadly categorized into axes such as protein synthesis, amino acid metabolism, nitrogen clearance, bile handling, drug detoxification, lipid and cholesterol homeostasis, glycogen storage, and more.^[6]^ To achieve all these tasks, the liver “divides and conquers” by delegating roles into distinct regions or “zones” across each functional unit of the liver, called lobules. Each liver lobule lies between two blood circulatory hubs, the portal triad (consisting of the portal vein, biliary duct, and hepatic artery) and the central vein.^[7]^ The zone near the portal triad is rich in oxygen and can support metabolic activities like gluconeogenesis and urea synthesis, while the zone around the central vein is oxygen-poor and resorts to glycolysis and is the site of drug detoxification. The term “zonation” refers to this separation of activity into distinct spatial regions. For implanted tissue to be broadly applicable in the clinic and perform a spectrum of the functions typically performed by the native liver, engineered tissues must replicate this zonal morphology and function.

We and others have previously developed engineered implantable human liver tissue grafts with the future goal of supplementing organ function.^[8]^ In one formulation, engineered tissues are composed of liver primary human hepatocytes (HuHep), normal human dermal fibroblasts (NHDF), and human umbilical vein endothelial cells (HUVEC) encased within fibrin hydrogel. This tissue not only carries out critical human liver functions and transcriptionally resembles the human liver, but also expands in response to the mouse hosts’ chronic, progressive liver injury. ^[8]^ However, early studies focused on building the human liver tissue and thus only examined explanted tissues at a single time point (12 weeks) after implantation and characterized the expression of only a few hepatic functional proteins. The extent to which such tissues express a larger subset of hepatic functional markers and importantly also replicate the spatial zonal phenotype of the native liver remains unknown. To bring engineered tissues closer to the clinic, a more comprehensive characterization of morphology and metabolic phenotype is needed.

Here, the dynamics of metabolic phenotype within engineered human liver tissues were studied after implant into mice with chronic and progressive liver injury. To evaluate metabolic maturity, a time-course characterization of both RNA expression and protein localization within human liver grafts after implantation was performed. This work identifies phenotypes within the engineered tissues that resemble the zonation found in the native human liver. These findings represent an important milestone towards human translation of therapeutic engineered liver tissue.

## 2. Results

### 2.1. Time course characterization of engineered human liver tissue graft expansion

We first sought to rigorously characterize the expansion dynamics of engineered implantable liver tissue created from HuHep, NHDF, and HUVEC within fibrin hydrogel (**Figure 1A**).^[8]^ HuHep were aggregated with NHDF using aggrewells before suspension in fibrin hydrogel with HUVEC. Grafts were implanted ectopically into the gonadal fat pad of FRGN mice. This mouse model experiences progressive liver injury unless treated with nitisinone (NTBC).^[9]^ NTBC was therefore cycled on and off to trigger chronic host liver injury and regeneration, along with concomitant stimulation of ectopic human liver graft growth. After implantation, grafts were characterized at five sequential time points during expansion from week 1 to week 9.5 by RNA sequencing, histology, and host serology (**Figure 1A**, **Figure S1A**).

**Figure 1.**
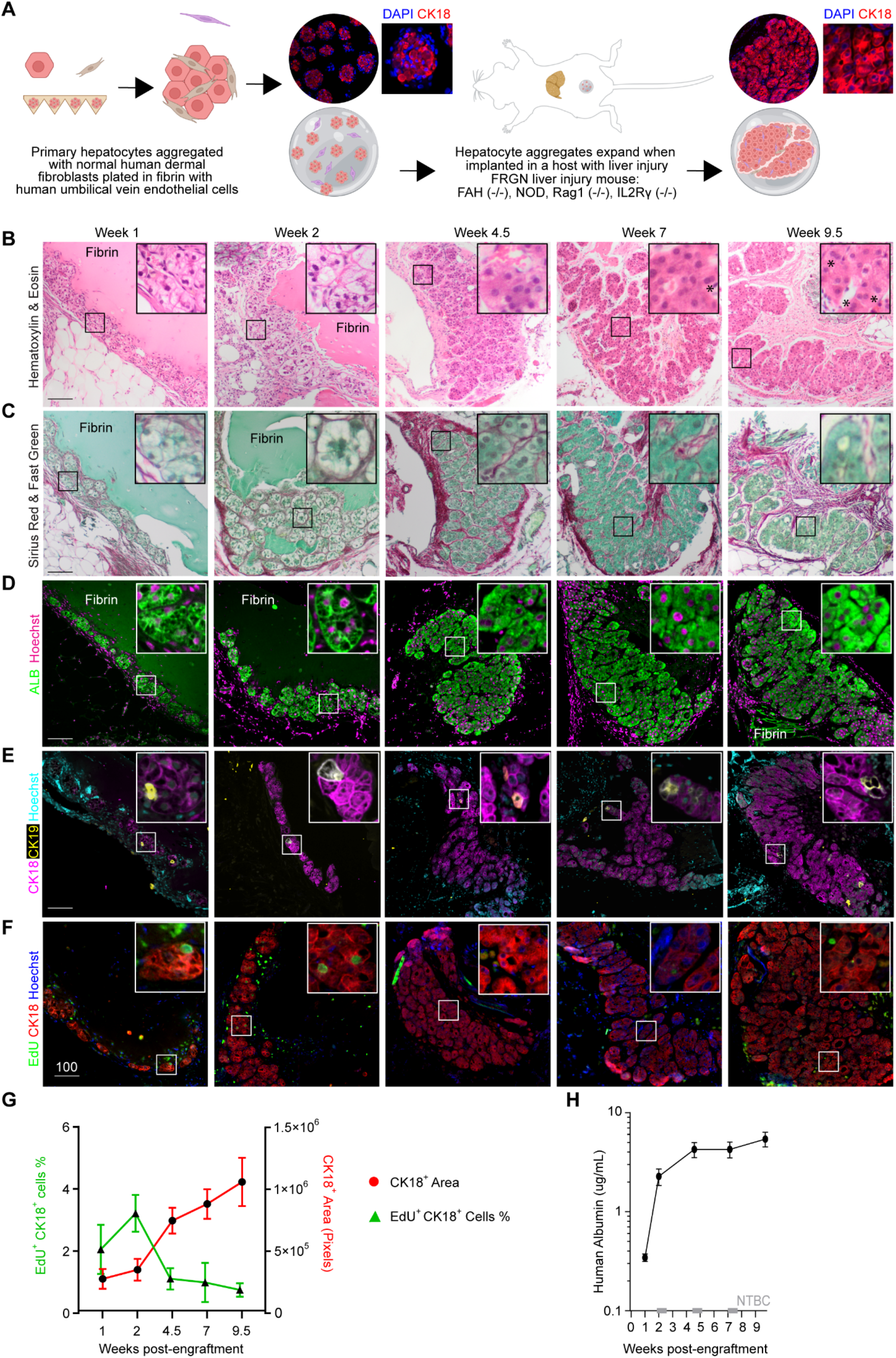
Time course characterization of engineered human liver tissue grafts. (**A**) Diagram depicting the construction of engineered human liver tissue grafts: primary human hepatocytes are aggregated with normal human dermal fibroblasts (NHDF) and then polymerized within fibrin with human umbilical vein endothelial cells (HUVEC). Grafts are implanted ectopically into mice with liver injury (FRGN mice. (**B**) Hematoxylin and eosin staining of engineered liver tissue grafts explanted at weeks 1-9.5. Nuclear content was stained with hematoxylin and basic compounds (proteins) were stained with eosin (right). >3 grafts from separate mouse hosts were stained for each panel at each time point; one representative graft is shown. Asterisk indicates binucleate cells. Scale bars display 100 μm. (**C**) Sirius red and fast green staining of engineered liver tissue grafts from weeks 1-9.5. Collagen was stained with Sirius red, and protein was counterstained with fast green (middle, right). >3 grafts from separate mouse hosts were stained for each panel at each time point; one representative graft is shown. Scale bars display 100 μm. (**D**) Immunostaining for hepatocyte marker albumin (ALB, green) and DNA (Hoechst, magenta) in engineered liver tissue grafts from weeks 1-9.5. >3 grafts from separate mouse hosts were stained for each panel at each time point; one representative graft is shown. Scale bars display 100 μm. (**E**) Immunostaining for hepatocyte marker CK18 (pink), biliary epithelial marker CK19 (yellow), and DNA (Hoechst, blue) in engineered liver tissue grafts from weeks 1-9.5. >3 grafts from separate mouse hosts were stained for each panel at each time point; one representative graft is shown. Scale bars display 100 μm. (**F**) Immunostaining for hepatocyte marker CK18 (red), EdU incorporation (green), and DNA (Hoechst, blue) in engineered liver tissue grafts grown from weeks 1-9.5. >3 grafts from separate mouse hosts were stained for each panel at each time point; one representative graft is shown. Scale bars display 100 μm. (**G**) The percentage of cells undergoing DNA replication, EdU+ nuclei (green, triangle), in the CK18+ area (red, circle), as quantified from the immunostaining performed in Figure 1F. Error bars represent the standard error of the mean. (**H**) Results from an ELISA for human albumin (μg/mL) from the blood of mice implanted with human liver grafts. Serum was collected at weeks 1, 2, 4.5, 7, and 9.5. NTBC was pulsed 3 times during graft growth and is shown in gray along the x-axis. Error bars represent the standard error of the mean.

To interrogate graft morphology over time post-implant, tissues were collected for immunostaining and histology (**Figure 1B-F**). Throughout the course of engraftment, engineered liver tissue self-organized to resemble the native human liver in epithelial structure (**Figure 1B**); 4.5 weeks after implantation, hepatocytes were densely packed and typically appeared as large cells with a low cytoplasmic to nuclear ratio.^[10]^ Some binucleated cells were observed (**Figure 1B**, asterisk). By weeks 7-9.5, engineered tissues were arranged into cord-like structures of cells encased by a collagen-rich matrix that resembled the native morphology of the human liver (**Figure 1C**).

Grafted cells expressed markers associated with hepatocytes, such as cytokeratin-18 (CK18) and the functional protein albumin (ALB), and these became more prominently expressed after 2 weeks of engraftment (**Figure 1D, F**). Rare, cytokeratin-19 (CK19)-positive cells, a marker of biliary epithelial cells and hepatic progenitor cells, were also observed (**Figure 1E**).

Quantification of the human hepatocyte area within the grafts (CK18+ area, red line, **Figure 1G**) demonstrated an increase in graft size over time. Quantification of 5-ethynyl-2’-deoxyuridine (EdU) incorporation revealed that there was an initial burst of hepatocytes undergoing DNA replication between weeks 1 and 2 after implant, followed by a drop-off in DNA replication as grafts continue to expand, possibly by cellular hypertrophy.^[11]^ Notably, hepatocyte cytoplasm appeared eosinophobic (likely protein-sparse) during the DNA replication phase at weeks 1-2, with cytoplasmic content recovering and appearing more eosinophilic (protein-dense) by week 4.5 (**Figure 1B, C**).

To monitor liver function, host blood was collected for serological assessment of human liver proteins by enzyme-linked immunosorbent assay (ELISA, **Figure 1G**). The secreted liver protein, albumin, was detected at each time point and its concentration in host blood increased over time, indicating increased liver function accompanying graft expansion. The most rapid rise in albumin production occurred between 1- and 2-weeks post-engraftment, followed by sustained protein expression.

### 2.2. Engineered human liver tissue shows increased metabolic capacity over time

To investigate the transcriptome of engineered liver tissues during expansion, bulk RNA was collected for RNA sequencing (RNAseq) in triplicate at weeks 1 - 9.5 after implant (**Figure S1A, B**). The number of reads in the bulk RNA samples showed increasing content mapping to the human genome over time (**Figure S1C**), indicating that samples became more enriched in human gene expression with graft growth, with fewer transcripts mapping to the murine hosts’ genome.

We then sought to further characterize the expression of genes associated with liver-specific functions throughout graft development. To do this, the expression of genes belonging to a hepatic signature in the engineered livers was visualized by a heatmap (**Figure 2A**). Most samples from later time points (10 out of 12 samples) after engraftment clustered together and separated from week 1, indicating a different transcriptional landscape after grafts are established in vivo. Two samples, one from week 4.5 and one from week 9.5, regularly clustered with the week 1 samples due to the high expression of fibrinogen genes (*FGB, FGA, FGG*) and serum amyloid genes (*SAA1, SAA2*) which were also highly expressed at week 1 (**Figure 2A, Figure S1B**).

**Figure 2.**
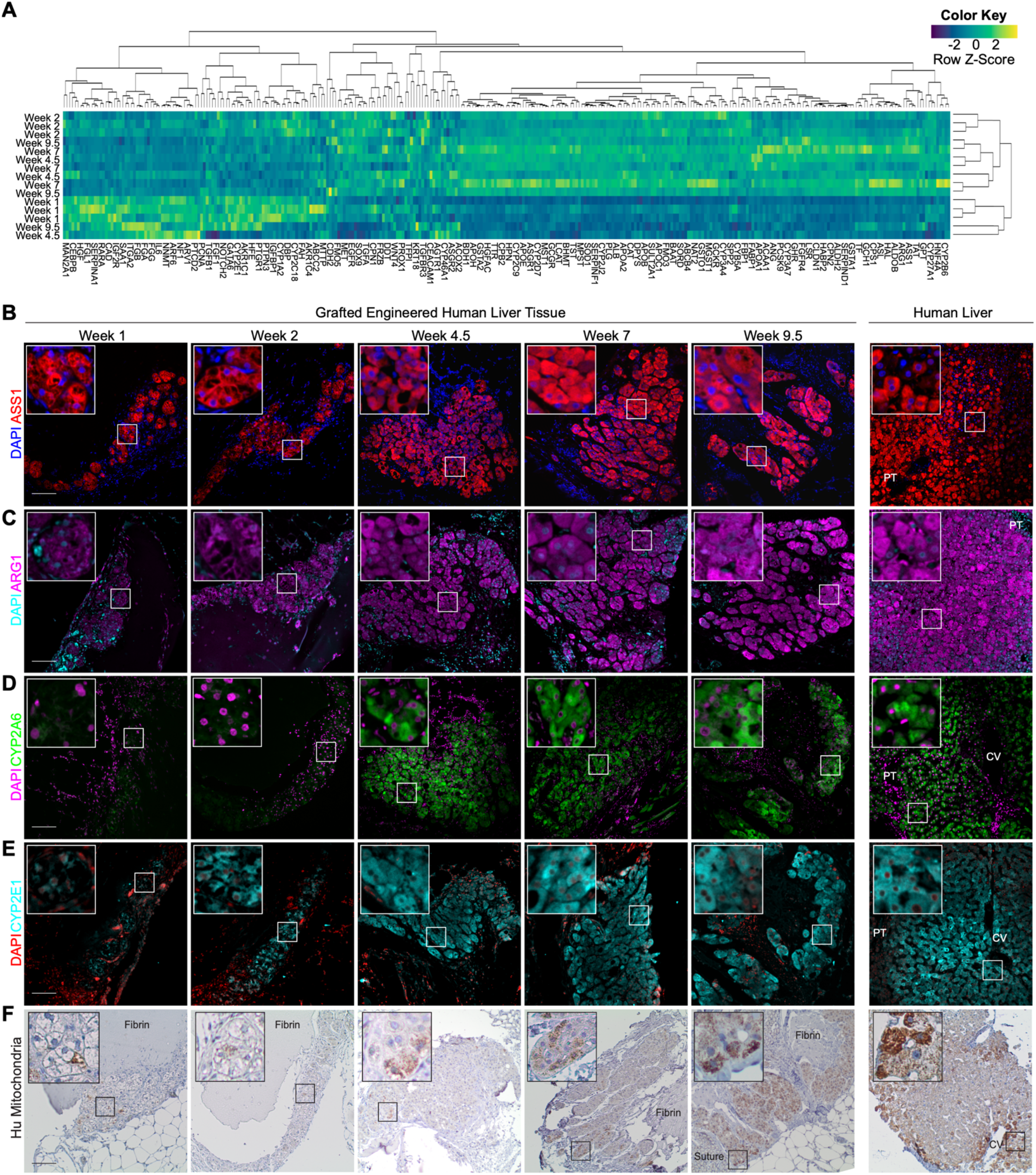
Engineered human liver tissue shows increased metabolic capacity over time. (**A**) Heatmap of liver-specific gene expression. Bulk RNA was collected and sequenced in triplicate from implanted grafts grown to weeks 1, 2, 4.5, 7, and 9.5 in separate mice. A heatmap plot was generated from transcripts per million of 276 genes from the liver gene set enrichment lists SU_LIVER (M7054) and HSIAO_LIVER_SPECIFIC_GENES (M13283).^[21,22]^ Corresponding Z-scores are shown for each row of genes. Hierarchical clustering was performed by Pearson correlation on rows and by Spearman for columns. (**B**) Immunostaining for ASS1 (red) and DNA (DAPI, blue) in engineered liver tissue grafts from weeks 1-9.5. >3 grafts from separate mouse hosts were stained for each panel at each time point; one representative graft is shown. Scale bars display 100 μm. PT = portal triad. (**C**) Immunostaining for ARG1 (pink) and DNA (DAPI, cyan) in engineered liver tissue grafts from weeks 1-9.5. >3 grafts from separate mouse hosts were stained for each panel at each time point; one representative graft is shown. Scale bars display 100 μm. PT = portal triad. (**D**) Immunostaining for CYP2A6 (green) and DNA (DAPI, pink) in engineered liver tissue grafts from weeks 1-9.5. >3 grafts from separate mouse hosts were stained for each panel at each time point; one representative graft is shown. Scale bars display 100 μm. CV = central vein, PT = portal triad. (**E**) Immunostaining for CYP2E1 (cyan) and DNA (DAPI, red) in engineered liver tissue grafts from weeks 1-9.5. >3 grafts from separate mouse hosts were stained for each panel at each time point; one representative graft is shown. Scale bars display 100 μm. CV = central vein, PT = portal triad. (**F**) Immunostaining for Hu-mitochondria (brown) and DNA (hematoxylin, blue) in engineered liver tissue grafts from weeks 1-9.5. >3 grafts from separate mouse hosts were stained for each panel at each time point; one representative graft is shown. Scale bars display 100 μm. CV = central vein.

We observed that genes commonly associated with liver regeneration such as *HGF* and *CCND1* were expressed early after implantation, along with *ARF6* and *GATA6* which are important in hepatogenesis (**Figure 2A**). ^[12,13]^ Notably, the course of early expression of such genes associated with liver regeneration largely matched the course of Ki67 protein expression by hepatocytes, further supporting the notion that at least some graft expansion was driven by early hepatocyte replication (**Figure 1G**).

Importantly, we further noted that a multitude of genes responsible for diverse hepatic functions such as *MPST, ALB, ASS1, GSTA2, ASGR1, CPS1, TTR*, and multiple cytochrome p450 genes showed increased expression over time after tissue implantation. This high metabolic gene signature was also observed by gene set enrichment analysis using iPathwayGuide (**Figure S2**). Thus, sequencing data suggested that genes associated with metabolic activities become upregulated with increased engraftment time in vivo.

To further characterize the metabolic maturation of grafted engineered liver tissues over time after implantation, we next immunostained graft sections for various functional proteins that represent key “axes”, or categories, of liver functions beyond protein production (**Figure 1D, H**), including xenobiotic metabolism, amino acid metabolism, and mitochondrial content (**Figure 2B-F**).^[14]^

One of the metabolic functional axes of the liver is amino acid metabolism and subsequent clearance of excess ammonia in the form of urea via the ornithine (urea) cycle. Argininosuccinate synthase 1 (ASS1, **Figure 2B**, red) is an enzyme that functions in the third step of this process to combine the amino acids citrulline and aspartate to form argininosuccinic acid, a precursor to arginine.^[15]^ Arginine is then hydrolyzed during the final step by arginase (ARG1, **Figure 2C**, pink) to form urea, for ammonia detoxification, and to form ornithine which is used for cell proliferation and collagen formation.^[16]^ Both ASS1 and ARG1 were expressed by engineered liver tissues and detectable throughout all growth stages.

Second, we sought to query the presence of proteins associated with xenobiotic (e.g., drugs and alcohol) detoxification, which is partly carried out by sequential cytochrome p450 enzyme processes. First, we interrogated CYP2A6, a cytochrome p450 enzyme involved in the metabolism of ~ 3% of drugs.^[17]^ Expression of CYP2A6 was not detected at weeks 1 and 2 but was observed at week 4.5 onward (**Figure 2D**, green). This is notable because modeling human CYP2A6 in vivo has remained challenging, as rodents exhibit species-specific variability in CYP2A6 activity.^[18]^ CYP2E1, a cytochrome p450 enzyme that catalyzes the oxidation of endobiotics, such as retinoids, is commonly studied in alcohol detoxification. Like CYP2A6 expression, we found that CYP2E1 protein expression was low at week 1 and became more prominent over time after engraftment (**Figure 2E**, cyan). However, CYP2E1 was detected at week 2 while CYP2A6 was not.

Finally, mitochondria are key metabolic hubs for hepatocyte homeostasis, flexibility, and survival.^[19]^ Some cytochrome p450 enzymes are also localized in the mitochondria.^[20]^ We found that the amount of mitochondrial content increased within human engineered liver tissue with increasing time of engraftment (**Figure 2F**, brown), like observations for CYP2E1 and CYP2A6.

Collectively, these data indicate RNA and protein expression of genes associated with hepatocyte identity and function arise in engineered human livers between weeks 1 and 2, followed by increased expression of several key metabolic functional proteins with increasing time following engraftment.

### 2.3. Engineered human liver tissues show a zonated phenotype

When exploring the expression of metabolic proteins within engineered liver grafts, we observed that some proteins, such as CYP2A6 and CYP2E1, appeared to be expressed heterogeneously across the hepatic graft (**Figure 2D, E**). We were intrigued by this, as in the native liver many functional genes and proteins are often “zonated”, such that their expression varies spatially across the liver lobule (**Figure 3A**).

**Figure 3.**
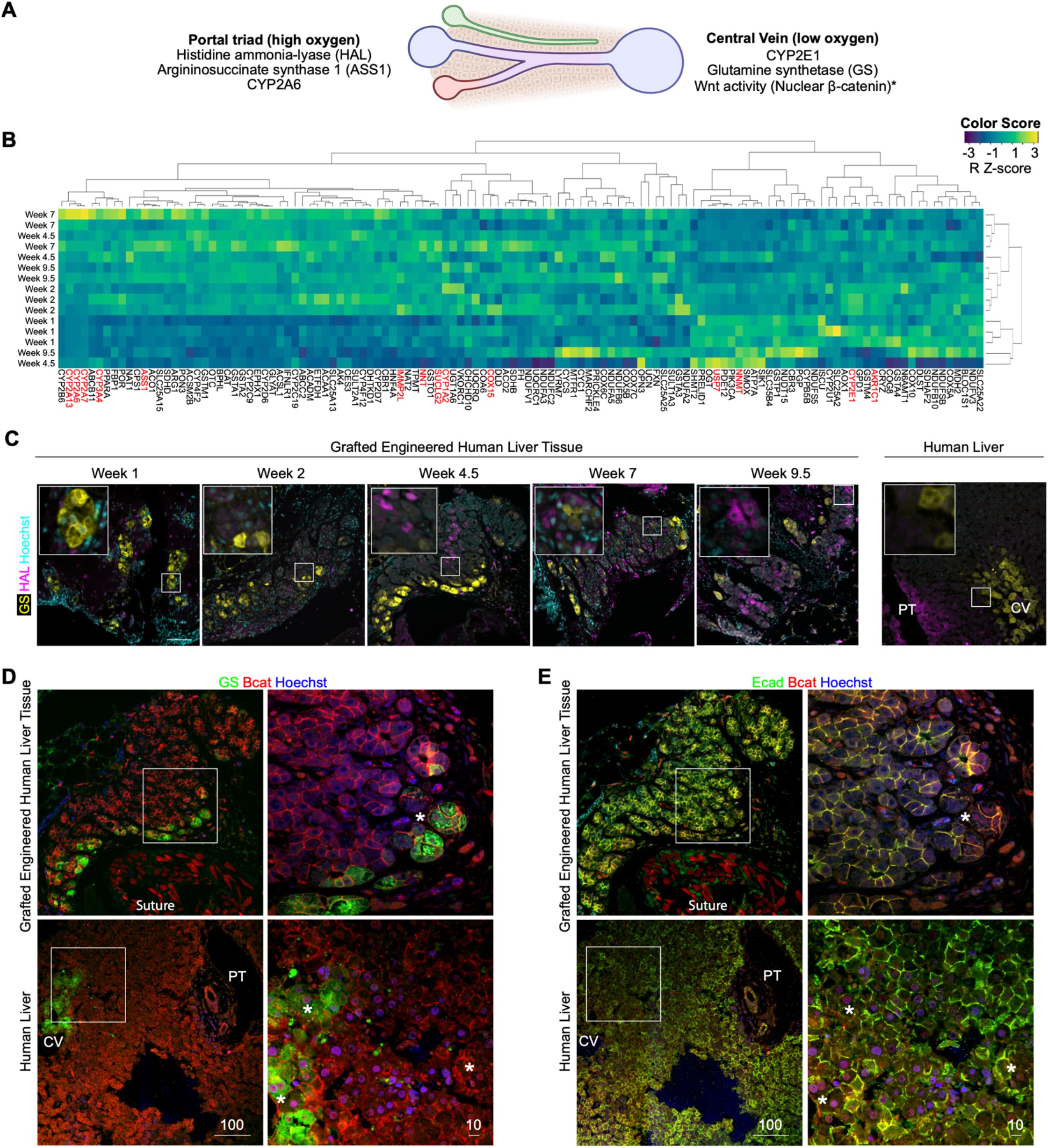
Engineered human liver tissues show a zonated phenotype. (**A**) Diagram of the liver lobule with corresponding zonated protein expression found at the portal triad or central vein. * = Reported in rodents. ^[23,24,25]^ (**B**) Heatmap of metabolic genes from zones 1, 2, and 3 of the liver. Bulk RNA was collected and sequenced in triplicate from implanted grafts grown to weeks 1, 2, 4.5, 7, and 9.5. A heatmap plot was generated from transcripts per million of genes related to liver metabolism. Corresponding Z-scores are shown for each row of genes. Hierarchical clustering was performed by Pearson correlation on rows and by Spearman for columns. The gene list was curated from the Gene Ontology (GO) terms: xenobiotic metabolic process, catabolic process, steroid metabolic process, glycoprotein metabolic process, lipoprotein metabolic process, carbohydrate metabolic process, cholesterol metabolic process, cellular respiration (oxidative metabolic process, oxidative metabolism, respiration), regulation of gluconeogenesis, urea metabolic process, glutamine synthesis, positive regulation of lipid biosynthetic process. Terms were chosen based on pathways that are functional at the portal triad and central vein of the liver lobule. Gene names highlighted in red appear zonated on the Human Protein Atlas (proteinatlas.org). (**C**) Immunostaining for GS (yellow), HAL (pink), and DNA (Hoechst, cyan) in engineered liver tissue grafts from weeks 1-9.5. >3 grafts from separate mouse hosts were stained for each panel at each time point; one representative graft is shown. Scale bars display 100 μm. CV = central vein, PT = portal triad. (**D**) Immunostaining for GS (green), β-catenin (red), and DNA (Hoechst, blue) in engineered liver tissue grafts grown for 4.5 weeks (top) and native human liver (bottom). >3 grafts from separate mouse hosts were stained for each panel at each time point; one representative graft is shown. Scale bars display 100 μm. CV = central vein, PT = portal triad. Asterisk indicates nuclear localization. CV = central vein, PT = portal triad. Inset scale bars display 10 μm. (**E**) Immunostaining for E-cadherin (Ecad, green), β-catenin (red), and DNA (Hoechst, blue) in engineered liver tissue grafts grown for 4.5 weeks (top) and native human liver (bottom). >3 grafts from separate mouse hosts were stained for each panel at each time point; one representative graft is shown. Scale bars display 100 μm. CV = central vein, PT = portal triad. Asterisk indicated nuclear localization. CV = central vein, PT = portal triad. Inset scale bars display 10 μm.

Thus, we next further characterized if engineered transplantable human liver grafts recapitulate aspects of metabolic zonation of native human liver. First, we further examined the expression of genes belonging to metabolic pathways that have zonated activity (**Figure 3B**), with genes that exhibit zonated protein expression marked in red in Figure 3B. We found that a majority of zonated human liver genes were expressed by week 2 post-implant by engineered human liver tissues.

To validate these findings, we next explored proteins that exhibit strict demarcation in their zonated expression in the human liver. First, we immunostained implanted engineered tissue grafts for histidase (HAL, **Figure 4C**, pink), which is responsible for the catabolism of histidine to transurocanic acid and ammonia and is expressed around the portal triad in the human liver. We found that HAL was restricted to the graft:host interface in engineered livers **(Figure 4C**). Glutamine synthetase (GS, **Figure 4C**, yellow) is important in nitrogen metabolism to synthesize glutamine from glutamate and ammonia. In the human liver, GS is only expressed in the first 1-3 cell layers surrounding the central vein. This was also consistent in grafts where only the first few cells inward from the graft periphery were positive for GS.

**Figure 4.**
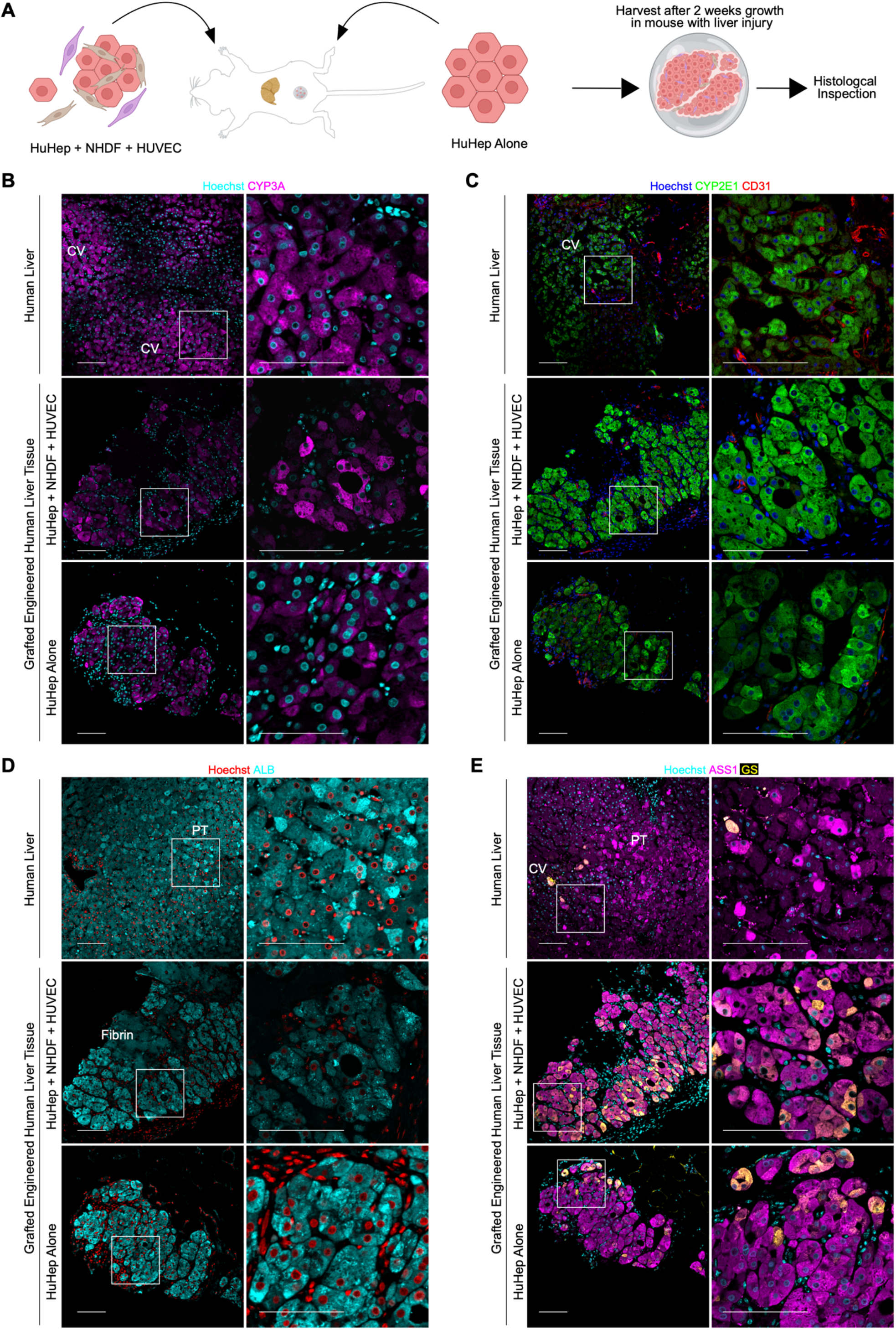
Human non-parenchymal supportive cells are not required for the zonated phenotype development in engineered liver tissue. (**A**) Diagram of experiment design: Grafts were generated from HuHep aggregates with and without nonparenchymal supportive cells NHDF and HUVEC. After two weeks of growth in hosts with liver injury, grafts were collected and stained for histological inspection. (**B**) Immunostaining for CYP3A (pink) and DNA (Hoechst, cyan) in human liver tissue (top), and engineered liver tissue grafts generated with NPC (middle) and without NPC (bottom) and grown in vivo for 2 weeks. >3 grafts from separate mouse hosts were stained for each panel; one representative graft is shown. Scale bars display 100 μm. CV = central vein. (**C**) Immunostaining for CYP2E1 (green), endothelial CD31 (red), and DNA (Hoechst, blue) in human liver tissue (top), and engineered liver tissue grafts generated with NPC (middle) and without NPC (bottom) and grown in vivo for 2 weeks. >3 grafts from separate mouse hosts were stained for each panel; one representative graft is shown. Scale bars display 100 μm. CV = central vein. (**D**) Immunostaining for ALB (cyan) and DNA (Hoechst, red) in human liver tissue (top), and engineered liver tissue grafts generated with NPC (middle) and without NPC (bottom) and grown in vivo for 2 weeks. >3 grafts from separate mouse hosts were stained for each panel; one representative graft is shown. Scale bars display 100 μm. PT = portal triad. (**E**) Immunostaining for ASS1 (pink), GS (yellow), and DNA (Hoechst, blue) in human liver tissue (top), and engineered liver tissue grafts generated with NPC (middle) and without NPC (bottom) and grown in vivo for 2 weeks. >3 grafts from separate mouse hosts were stained for each panel; one representative graft is shown. Scale bars display 100 μm. CV = central vein, PT = portal triad.

In rodents, β-catenin is required for zonation and expression of GS, and its inhibition results in a periportal phenotype. ^[23,24,25]^ The expression and localization of β-catenin were assessed by immunostaining in both human liver and engineered liver tissue (**Figure 4D**). In the absence of the Wnt ligand, β-catenin will either be sequestered at the membrane and associated with E-cadherin or in the cytoplasm degraded by the APC destruction complex.^[26]^ In the presence of Wnt, β-catenin will become stabilized and enter the nucleus to induce the expression of Wnt genes, such as *GS*. In human liver tissue, nuclear β-catenin was observed at and around the central vein, as well as throughout the liver lobule (**Figure 4D**, upper). In engineered human liver tissue, nuclear β-catenin was difficult to observe but was detectable in regions that were co-expressing GS (**Figure 4D**, lower, asterisk). When β-catenin was not observed in the nucleus it was localized at the membrane near E-cadherin (**Figure 4E**).

Finally, we sought to characterize the spatial expression of a protein that is zonated differently between species, in this case, humans *vs*. mice. We immunostained for E-cadherin protein, which is zonated in mice but not in the human liver.^[28]^ We found that like human liver tissue, E-cadherin was expressed uniformly (i.e., not zonated) across engrafted engineered liver tissues. Thus, engineered human liver tissue could serve as an important model system that recapitulates various aspects of human metabolic zonation.

### 2.4. Human non-parenchymal supportive cells are not required for the zonated phenotype development in engineered liver tissue

Rodent liver zonation is directed by oxygen gradients, Wnt molecules, and other signals originating in part from nonparenchymal cells (NPC) in the native liver, such as central vein endothelial cells.^[29]^ We thus wondered the extent to which expression of metabolic markers and zonation of engineered human liver tissues was dependent upon the inclusion of the HUVECs and NHDFs in the engineered tissue. We set out to further characterize implanted engineered liver tissue grafts fabricated with and without NPC (NHDF or HUVEC, **Figure 4A**).

We fabricated engineered liver tissues with and without NPC and engrafted these tissues into FRGN mice. After two weeks of engraftment, both engineered liver tissues consisting of hepatocytes alone and hepatocytes with NPC expressed proteins that are important for metabolic function and zonation, such as cytochrome p450 enzyme CYP3A (**Figure 4B**), CYP2E1 (**Figure 4C**), albumin (**Figure 4D**), ASS1 (**Figure 4E**) and GS (**Figure 4E**). CD31 is expressed by endothelial cells and was observed in grafts with and without NPC (**Figure 5C**). In the grafts that were generated without HUVEC NPC, the CD31-positive cells are most likely of host origin, and it has been previously reported that host circulation connects to grafts after implant and ultimately becomes chimeric.^[8]^

Notably, GS expression was detected at the graft periphery in a zonated pattern in both graft formulations. This suggests that, while supportive factors from HUVEC or NHDF are important to promote graft expansion,^[8]^ they are not required to induce the expression of zonated proteins in human hepatocytes in engineered human livers. We noted that in both tissues with and without NPC, GS tended to be localized to the graft border, near the interface between the graft and host tissues. The drivers of the zonated phenotype may be originating from the adjacent host tissue or the host circulation.

## 3. Conclusion

The present study used RNAseq, immunostaining, and host serology to characterize the molecular landscape and metabolic maturation of engineered human liver tissue expansion. After implant in mice experiencing liver injury, engineered liver grafts expressed genes and proteins that are important in liver metabolic function. In the native liver, many metabolic functions are spatially delegated to different populations of hepatocytes depending on their proximity to the major blood vessels, a phenomenon termed liver zonation. The expression of proteins that are zonated in the native liver also appeared to be zonated in the engineered liver. This report not only brings these tissues closer to relevance for the clinic but also offers the field a platform to study the mediators that establish human liver zonation.

In rodent livers that are expanding after surgical resection (hepatectomy), there is a reduction in glycogen content, a decrease in serum glucose, less metabolic pathway gene expression,^[30]^ and a switch to oxidation to produce metabolites and energy.^[31]^ This metabolic stress results in changes in the redox metabolism that can promote regeneration and transition cells from quiescence to proliferation.^[32]^ The transition to proliferation corresponds with the downregulation of amino acid biosynthesis,^[30]^ which is reminiscent of the eosinophobic cytoplasms observed during DNA replication at weeks 1 and 2 of engineered graft expansion. Similarly, we also noted an increase in metabolic protein expression after replication had lessened.

The cytochrome p450 enzymes CYP2E1 and CYP2A6 were not detected at early time points after engineered liver tissue engraftment but were expressed by week 4.5. This may suggest that either xenobiotic metabolism was not required until this point, or the engineered livers didn’t have the capacity for xenobiotic metabolism until cell division had lessened. This agrees with observations in rodent models of liver regeneration where CYP enzyme expression is reduced while cells are dividing. ^[33,34,35]^ Similarly, mitochondria are an important component of metabolic function, but mitochondrial content was very low at early time points during cell division when organelles are disrupted. These observations are also mirrored in rodent liver regeneration, wherein hepatocyte metabolic activity is reduced in the early stages of liver regeneration, and metabolic potential increases steadily over time thereafter.^[30]^

Some factors that regulate liver zonation include the availability of oxygen across the lobule and the Wnt pathway.^[23–27]^ In rodents, Wnt ligands are secreted by liver central vein endothelial cells to regulate β-catenin and induce GS expression.^[24,25]^ Based on this, we initially hypothesized that the HUVEC nonparenchymal cells within the engineered liver tissues would not only support hepatocyte growth, as has been previously shown,^[8]^ but also support a zonation phenotype. Surprisingly, the engineered livers showed a zonated phenotype in the absence of human NPC from graft formulation. Wnt proteins are not traditionally described as circulating to distant sites or acting as hormones, which makes it unlikely that Wnts produced by the host’s central vein endothelial cells would affect the ectopic graft.

We further observed that pericentral proteins such as GS were largely expressed in spatial positions adjacent to host tissue, suggesting that interactions with the host tissue impacted the emergence of human liver tissue zonation. It is possible that Wnts are supplied by endothelial cells in the fat pad of the host and direct a zonated phenotype in the graft. Alternatively, this conundrum may support the notion that circulatory factors, such as nutrients or high ammonia levels, are partly responsible for the expression of zonated proteins such as GS, ASS1, ARG1, and HAL. In this case, Wnt signaling may thus be a lesser contributor to the re-establishment of zonation in engineered human liver tissue generated from *adult* hepatocytes. The mechanisms for Wnt transport and contribution to zonation in the human setting, at least in engrafted human liver tissues, are yet to be unveiled.

Intensive engineered devices that establish gradients of nutrients or oxygen exposure are necessary to study human zonation in vitro. ^[35–47]^ To our knowledge, this is the first report of an engineered human-derived zonated phenotype in vivo. Future studies could leverage this system to uncover mechanisms of both human hepatocyte expansion and zonation in vivo or to create zonated human liver tissues for the treatment of liver disease.

## 4. Experimental Section/Methods

### Cell sources

Cryopreserved primary human hepatocytes were purchased from Thermo Fisher Scientific, lot hu1880 (34-year-old, white, female donor) was used in **Figure 1 – Figure 3**, lot hu8375 (19-year-old, white, female donor) was used for **Figure 4**. Primary human umbilical endothelial cells (HUVEC) and normal human dermal fibroblasts (NHDF) were purchased from Lonza and used at passage < 8. HUVEC were maintained in dishes in EGM-2 media (Lonza), and NHDF were maintained in DMEM (Thermofisher Scientific) with 10% fetal bovine serum (FBS, Gibco) and 1% penicillin-streptomycin (penstrep, VWR).

### Aggregate culture

Aggregates were generated as previously described. ^[5,45]^ Briefly, human primary hepatocytes were thawed and immediately plated into pluronic-coated AggreWell micromolds at 600,000 cells per well on a 6-well plate. Aggregates of HuHep with NHDF were generated by adding 1 million NHDF cells per well at the time of plating. Cells were incubated overnight to gravity-settle and form aggregates in high glucose DMEM with 10% FBS, 1% ITS supplement (insulin, transferrin, sodium selenite; BD Biosciences), 0.49 pg/mL glucagon, 0.08 ng/mL dexamethasone, 0.018 M HEPES, and 1% pen strep.

### Graft construction

To create hepatic aggregates, human primary hepatocytes were thawed and immediately plated into AggreWell micromolds along with NHDF (approximately 100 hepatocytes and 160 dermal fibroblasts per aggregate) and incubated overnight.^[48]^ HUVEC and hepatic aggregates were then suspended into 10 mg/ml fibrin at a concentration of 1,250,000 HUVEC/mL and 45,000 aggregates/mL and allowed to polymerize within a 16 mm diameter polydimethylsulfoxide circular mold. Engineered tissues were cut with a 6 mm biopsy punch immediately before implantation. Each engineered tissue contained approximately 450,000 human hepatocytes, 720,000 dermal fibroblasts, and 125,000 HUVECs upon implantation.

### Graft implantation

All experiments were performed in female FAH-/-, RAG2-/-, IL2RG-/- NOD (FRGN) mice, an immune-deficient model of hereditary tyrosinemia type I. Mice were anesthetized using isoflurane, and bioartificial tissues were implanted onto the perigonadal fat pads. Three 6 mm tissues per mouse were ligated to the fat by passing a 5-0 suture through both the construct and the fat. Surgical incisions were closed aseptically, and mice were administered 1 mg/kg buprenorphine (slow releasing). Nitisinone (NTBC) was withdrawn from animals’ drinking water immediately after implantation.

### Mouse model of liver injury

The FAH-/-, RAG2-/-, IL2RG-/-, NOD (FRGN) mouse strain is an immune-deficient model of hereditary tyrosinemia type I, an established model of chronic liver injury. These mice experience progressive liver failure unless the small molecule nitisinone (NTBC) is administered in the animal’s drinking water. Upon artificial tissue implantation, NTBC was cycled on/off NTBC to induce moderate liver injury using established protocols.^[8]^

### Number of animals studied

Power calculations in which one tissue is implanted into a given mouse assume an alpha (p-value) of 0.05 and a power of 0.90. Based on our previous studies, a sample size of 8 is adequate to detect statistically significant differences in liver injury studies. To allow for mortality in liver injury studies in which animals are cycled on/off NTBC (typically 20% mortality), an additional two animals are enrolled per group to bring the total to 10 animals per group per experiment. The number of animals per endpoint at experiment set up is shown in Figure S1.

### RNA sequencing

Tissues were excised from animals and placed in RNALater (Thermo Fisher) to stabilize RNA. Tissues were manually dissected to remove excess mouse fat from around the implanted tissue and prevent the overrepresentation of murine transcripts from the downstream sequencing libraries. A phenol-chloroform extraction protocol was used to isolate RNA from tissues. RNA concentration was measured through NanoDrop and frozen. Frozen RNA was sent to BGI Genomics (Cambridge, Massachusetts, U.S.A.) for sequencing. Standard RNAseq analysis was performed. Reads were aligned to a human reference genome and the number of reads mapped to individual genes and transcripts was counted. Heatmaps to visualize gene expression from RNAseq analysis were generated from the normalized transcripts per million for each gene using R and RStudio gplots and pheatmap packages. Hierarchical clustering was performed by Pearson correlation on rows and by Spearman for columns; z scores are reported.

### Gene Set Enrichment Analysis

The significantly impacted pathways and biological processes were analyzed using iPathwayGuide (Advaita Bioinformatics).^[49,50]^ Lists of differentially expressed genes produced during RNAseq analysis were input into the meta-analysis program. Only the following comparisons produced enough DEGs for analysis: week 2 vs. week 1, week 4.5 vs. week 1, week 7 vs. week 1, week 9.5 vs. week 1, and week 7 vs. week 2.

### ELISA

Throughout the experiment and at the time of sacrifice, mice were bled through the saphenous vein, and blood was centrifuged to isolate the serum. Human albumin protein was quantified within mouse serum by enzyme-linked immunosorbent assay (ELISA) using goat polyclonal capture and HRP-conjugated goat antihuman albumin antibodies (Bethyl E80129). Plots were generated using GraphPad Prism.

### Immunostaining

Engineered tissues and host auxiliary tissues were harvested and formalin-fixed, paraffin-embedded for histology. Embedded samples were sectioned, and heat-induced antigen retrieval was performed in a pressure cooker with sodium citrate buffer. Samples were blocked with 1.5% normal goat serum, followed by incubation with primary antibodies listed in **Table 1** overnight at 4°C, and incubation with secondary antibodies for 1 hour at room temperature or overnight at 4°C. Secondary antibodies used were donkey anti-goat Alexa Fluor 488 (Invitrogen), donkey anti-goat IgG Alexa Fluor 488 conjugate (Invitrogen), donkey anti-mouse Alexa Fluor® 555 conjugate (Invitrogen), donkey anti-rabbit IgG Alexa Fluor 594 conjugate (Novex), and goat anti-mouse IgG1 Alexa Fluor 555 conjugate (Novex). Nuclei were counterstained with Hoechst or DAPI. Immunohistochemical detection of human mitochondria was performed using the mouse-specific HRP/DAB (ABC) Detection IHC Kit (ab64259). Images were acquired using the microscopes specified in the following section.

**Table 1.** List of antibodies used for immunostaining.

| Antibody Target | Supplier | Catalog | Host species | Dilution |
| --- | --- | --- | --- | --- |
| CK18 | Agilent | M701029-2 | Mouse | 1:100 |
| CK19 | Abcam | ab52625 | Rabbit | 1:250 |
| Albumin | Bethyl | A80-129A | Goat | 1:100 |
| VIM | Abcam | ab45939 | Rabbit | 1:1000 |
| CD31 | Agilent | M0823 | Mouse | 1:20 |
| ARG1 | Sigma | HPA003595 | Rabbit | 1:400 |
| CYP2A6 | Sigma | HPA047262 | Rabbit | 1:200 |
| ASS1 | Sigma | HPA020934 | Rabbit | 1:200 |
| HAL | Sigma | HPA038547 | Rabbit | 1:200 |
| GS | Millipore | MAB302 | Mouse | 1:100 |
| CYP2E1 | Abcam | ab28146 | Rabbit | 1:100 |
| Hu Mitochondria | Abcam | ab92824 | Mouse | 1:800 |
| ECAD | R&D | AF748 | Mouse | 1:100 |
| BCAT | Cell Signaling | mAb #8480 | Rabbit | 1:100 |
| CYP3A | Santa Cruz | sc-365415 | Mouse | 1:100 |

### Microscopy

Images were acquired using a Nikon wide-field epifluorescence microscope, Nikon A1R confocal, and Yokogawa W1 spinning disk confocal. All microscopes were run using Nikon Elements software. The widefield microscope was an inverted Nikon Eclipse TiE system with a 120BOOST LED-based illumination system and equipped with a Photometrics HQ2 CoolSnap camera and motorized XY stage. The Nikon A1R point scanning confocal system was run on an inverted Nikon Eclipse TiE base with 405-, 488-, 568- and 647nm excitation laser lines and four detectors: two GaAsP and two Alkali PMTs with a motorized XY stage. The Yokogawa W1 spinning disk confocal has an inverted Nikon Eclipse TiE base and 100mW 405-, 490-, 561-, and 640nm lasers, equipped with an Andor iXon 888 Life EMCCD camera linked with a 10-position filter wheel and a motorized XY stage. The spinning disk system was enclosed in an environmental chamber with temperature and local [CO2] control.

### EdU staining and quantification

Animals were injected with 50 mg/kg 5-ethynyl-2’deoxyuridine (EdU, Thermo Fisher) one hour before sacrifice. At the time of sacrifice, animal tissues (peritoneal implants and fat pads, liver, and small intestine) were excised, rinsed in PBS, and fixed in 4% PFA at 4°C for two days. After fixation, tissues were dehydrated to 70% ethanol, paraffin-embedded, and sectioned into 6 μm-thick sections for mounting onto histology slides. To visualize EdU incorporation in human hepatocytes undergoing DNA replication at the time of injection, slides were deparaffinized and co-stained with an Alexa Fluor 647 click chemistry conjugation kit (Invitrogen Click-iT™), mouse anti-human CK18, and Hoechst. Images were acquired using a wide-field epifluorescence microscope (listed above). To quantify percent positive hepatocyte nuclei, ilastik software (http://ilastik.org) was trained to recognize EdU-positive Hoechst-positive nuclei within CK18-positive cells. This training was performed on three images from different animal groups to ensure training diversity. ilastik’s Density Count feature was used to count the EdU-positive hepatocyte nuclei, and all images were batch processed to ensure they were counted identically. The total nucleus count was performed using the same process, but by training the software to recognize all Hoechst-positive nuclei within CK18-positive cells, not just the EdU-positive nuclei. The EdU-positive number was divided by the total nucleus count to obtain a percentage of total nuclei undergoing DNA replication at the time of EdU incorporation.

## Acknowledgments

This research was funded by NIH NIDDK (R01DK128551) (K.R.S.), the NCATS Translational Research Training Program TL1 TR002318 and the NIGMS Molecular Medicine Training Grant T32 GM095421 (C.L.F), the Environmental Pathology/Toxicology NIEHS training program T32 ES007032 and the NSF Postdoctoral Fellowship in Biology Award Number 2109919 (T.N.M.), NIH NIBIB Cardiovascular training grant (S.H.S., T32EB001650), and the University of Washington Post-Baccalaureate Research Education Program (C.B.A.). Any opinions, findings, conclusions, or recommendations expressed in this material are those of the author(s) and do not necessarily reflect the views of the funders. Figures were created with BioRender.com. We thank Dr. Dale Hailey from the UW Garvey Imaging Core for assistance with imaging. We thank Dr. Theo Bammler and Dr. Lu Wang of the Integrative Environmental Health Sciences Facility Core as part of the Interdisciplinary Center for Exposures, Diseases, Genomics, and Environment at UW. We thank all the fantastic animal husbandry and veterinary staff in The Department of Comparative Medicine’s vivarium. We thank all patient tissue donors for their generous contributions to science. We acknowledge that the experiments described in this text were conducted on the land of the past, present, and future Coast Salish peoples, the land that touches the shared waters of all tribes and bands within the Suquamish, Tulalip, and Muckleshoot nations; we honor the land itself and these Coast Salish tribes.

## Data Availability Statement

Data described in this manuscript are available upon request.

---

Successfully engineered human liver tissue should perform functions of and structurally resemble the native liver. This includes mimicking the characteristic of liver tissue known as “zonation”, where different roles are carried out in distinct regions. Here we describe a time-course evaluation of engineered human liver tissue implanted in rodents in vivo and demonstrate a zonated phenotype of metabolic proteins.

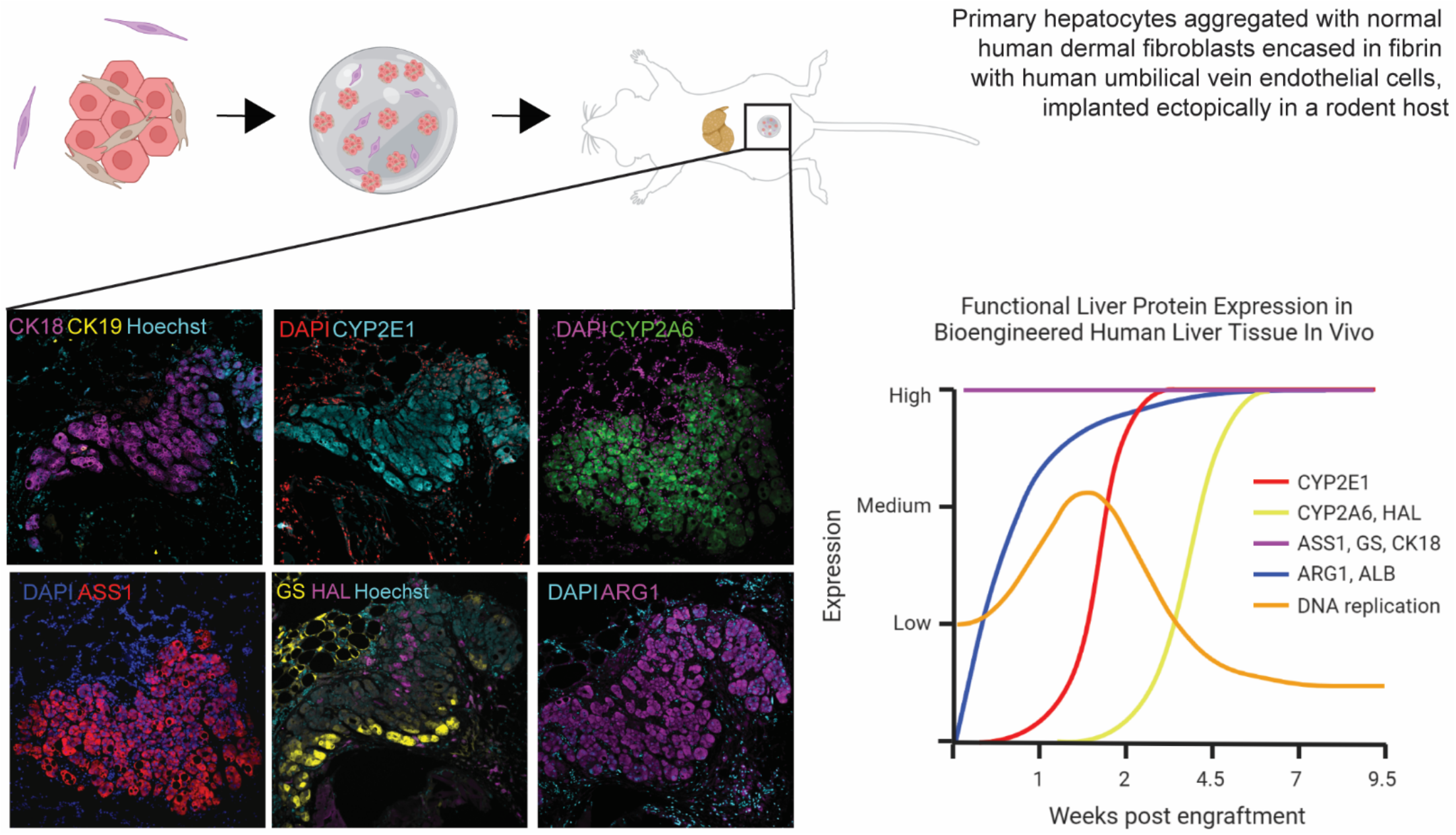

## Supporting Information

**Figure S 1.**
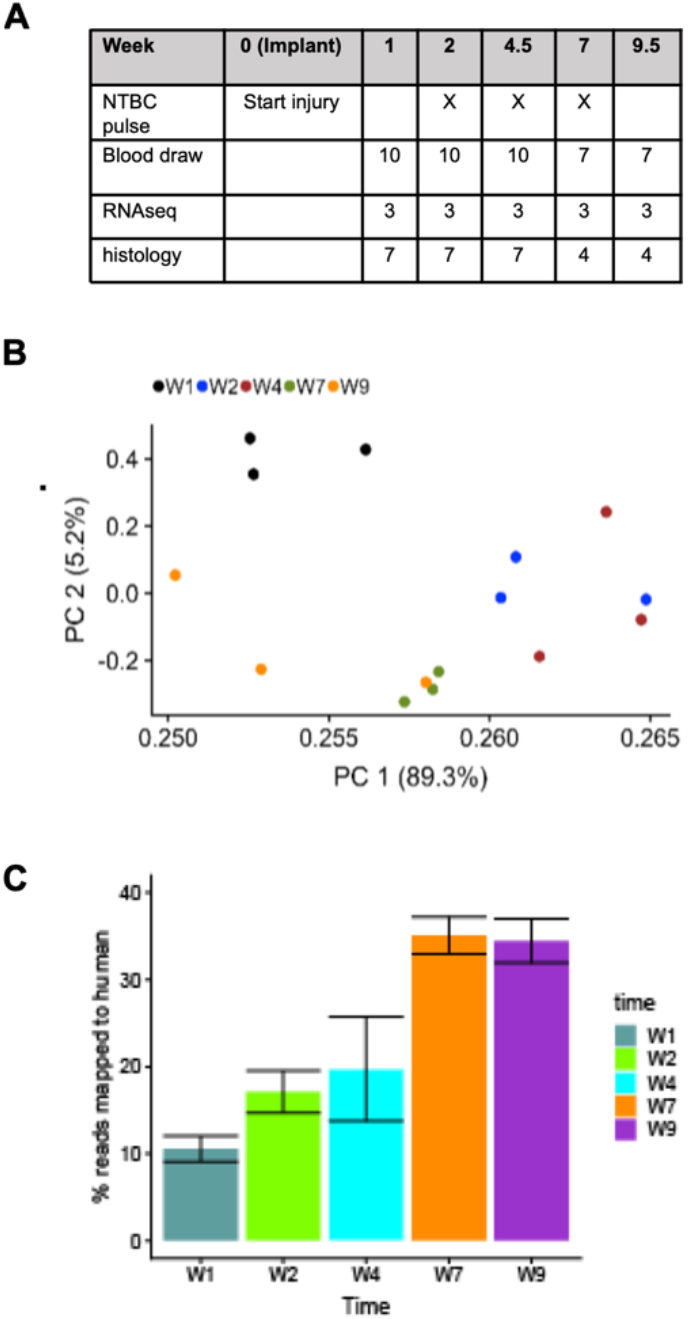
related to Figure 1. Time course characterization of engineered human liver tissue grafts. (**A**) Table of timeline, animal treatments (shown as “X”), and the number of engineered liver grafts collected from separate animal hosts for each endpoint (blood draw, RNAseq, histology). (**B**) Principal component plot for each sample of three replicates from RNA-sequencing. Three different grafts implanted in three different mouse hosts were collected and sequenced individually. Week 1 = W1, week 2 = W2, etc. Samples from week 4.5 (W4) were tightly clustered replicates, while replicate samples from week 9.5 (W9) had greater variability. (**C**) Bar chart showing the percentage of reads in the RNAseq data that were mapped to human origin (y-axis) at each time point (x-axis), as opposed to RNAseq reads that mapped to the mouse host. Week 1 = W1, week 2 = W2, etc. Replicate samples from separate grafts implanted in separate hosts are shown here; the bar chart shows the standard error of the mean.

**Figure S 2.**
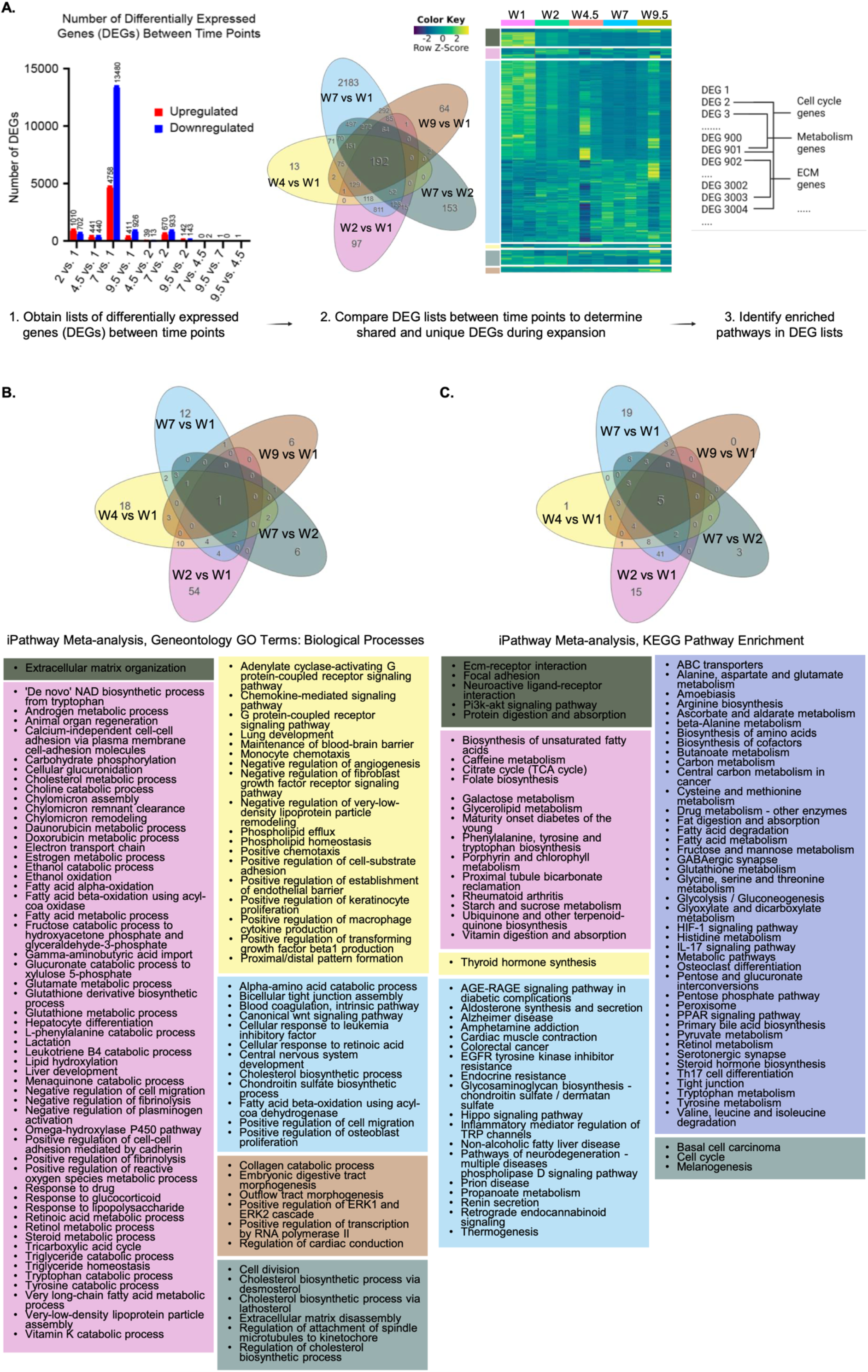
related to Figure 2. Hallmarks of engineered liver tissue development include extracellular matrix remodeling, expansion, and metabolic activation. (**A**) Diagram of workflow for differential expression analysis and gene set enrichment: Differentially expressed genes (DEGs) were identified by comparing each time point (left), number of DEGs is shown by bar chart, color represents upregulated (red) or downregulated (blue) genes. DEGs for the 5 most disparate comparisons were input into iPathwayGuide meta-analysis to determine unique and shared DEGs for each comparison – shown by the Venn diagram and heatmap (middle). Significant DEGs were then input into gene set enrichment analysis to explore GO terms and KEGG pathways. (**B**) Results from iPathwayGuide meta-analysis described in (A). Significant DEG list was input into iPathwayGuide metaanalysis, and the resulting Geneontology GO terms: Biological Processes that were enriched in the DEG list are shown. GO terms with an asterisk have been abbreviated (ex. G-protein-coupled receptor signaling pathway is GPCR signaling pathway). Prominent terms include those related to cell adhesion and the extracellular matrix, cell division, and metabolic processes. The color indicates which comparison the GO term belongs to, ex. pink is W2 vs. W1. (**C**) Results from iPathwayGuide meta-analysis described in (A). Significant DEG list was input into iPathwayGuide meta-analysis, and the resulting KEGG pathways that were enriched in the DEG list are shown. GO terms with an asterisk have been abbreviated (ex. G-protein-coupled receptor signaling pathway is GPCR signaling pathway). Prominent terms include those related to cell adhesion and the extracellular matrix, cell division, and metabolic processes. The color indicates which comparison the GO term belongs to, ex. pink is W2 vs. W1.

